# Rapid and accurate assembly of large DNA assisted by *in vitro* packaging of bacteriophage

**DOI:** 10.1101/2022.08.01.502418

**Authors:** Shingo Nozaki

## Abstract

Development of DNA assembly methods made it possible to construct large DNA. However, achieving the large DNA assembly easily, accurately, and at low cost remains a challenge. This study shows that DNA assembled only by annealing of overlapping single-stranded DNA ends, which are generated by exonuclease treatment, without ligation can be packaged in phage particles and can also be transduced into bacterial cells. Based on this, I developed a simple method to construct long DNA of about 40 - 50 kb from multiple PCR fragments using the bacteriophage *in vitro* packaging system. This method, named iPac (*<u>i</u>n vitro* <u>Pac</u>kaging-assisted DNA assembly), allowed accurate and rapid construction of large plasmids and phage genomes. This simple method will accelerate research in molecular and synthetic biology, including the construction of gene circuits or the engineering of metabolic pathways.

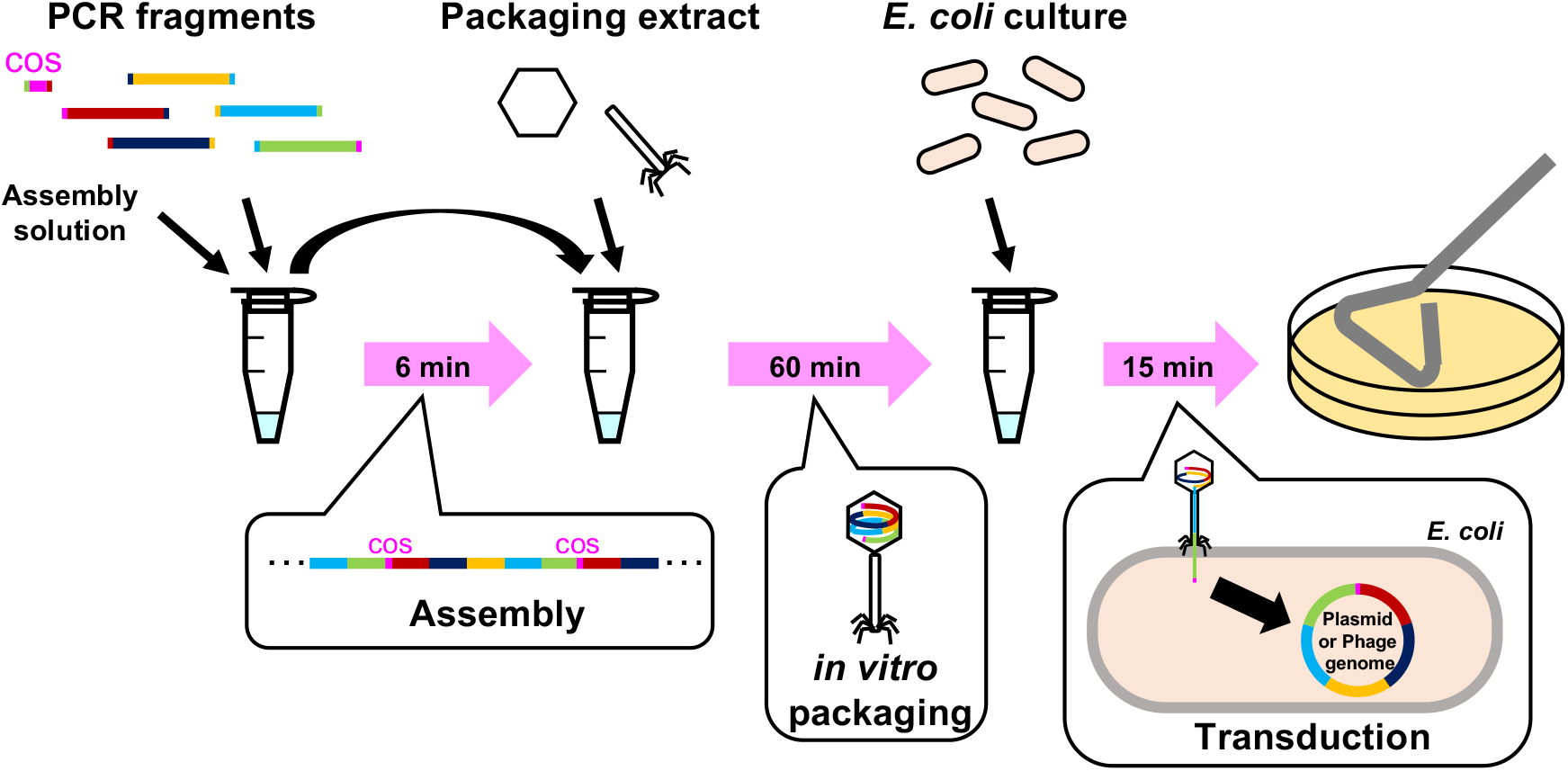

## Introduction

DNA assembly is one of the fundamental techniques in molecular and synthetic biology. The development of DNA assembly technology has made it possible to assemble various DNA fragments in the desired order. In recent years, there has been an increasing demand for construction larger and more complex DNA, such as constructing synthetic genomes ^1,2^. However, it is still difficult to achieve the large and complex DNA assembly accurately, efficiently, inexpensively, and easily.

DNA cloning into a plasmid vector, namely, two-fragment assembly, was first performed by using restriction enzyme and DNA ligase ^3^. This has led to the development of recombinant DNA technology. As a technique for cloning large DNA with restriction enzymes and DNA ligase, cloning into λ phage vector using *in vitro* packaging of λ phage have been developed ^4–7^. In DNA cloning using the λ phage vector, the λ phage genome itself is used as a vector. λ phage genome is composed of 3 domains: left and right arms, which are essential for lytic cycle including phage proliferation and infection to *Escherichia coli,* and replaceable region involved in lysogenic cycle between the left and right arms. In the λ phage vector cloning, the replaceable region is replaced with a DNA fragment to be cloned. The total size including the left and right arms and the cloned DNA should be less than 52 kb, which is the maximum size capable of the λ phage capsid, which can package DNA of 38 – 52 kb ^8^. The length of the left and right arms is about 29 kb, so about DNA fragment of 9 – 23 kb can be cloned with this system. To clone longer DNA, cosmid cloning was developed ^9^. The cosmid cloning uses plasmid vectors containing the cos sequence, which is required for the λ phage packaging system. The size of the cosmid vector is significantly smaller than the λ phage vector, and it is possible to clone a DNA fragment of about 30 – 45 kb. Recently, long regions such as natural product pathways have also been cloned by combining the *in vitro* packaging system of λ phage with the CRISPR-Cas9 system ^10^.

In recent years, for the construction of plasmids from multiple DNA fragments, seamless assembly techniques such as SLIC (Sequence and Ligation-Independent Cloning), In-Fusion, Gibson assembly and Golden Gate assembly are becoming mainstream ^11–15^. In the seamless assembly, the ends of DNA fragments to be assembled are designed to overlap each other. SLIC uses 3’-5’ exonuclease activity of T4 DNA polymerase to generate 5’ overhangs at the ends of insert(s) and a linearized vector. By annealing the overlapping sequences of about 25 bp, the insert(s) and the vector are assembled *in vitro.* In the same way, In-Fusion can assemble DNA fragments by using Vaccinia virus DNA polymerase with shorter DNA overlapping of about 15 bp. Gibson assembly uses thermostable DNA polymerase and DNA ligase in addition to thermolabile T5 5’-3’ DNA exonuclease. After insert(s) and a linearized vector DNA is mixed with these enzymes at 50°C, the exonuclease resects the ends of the DNA fragments to generate 3’ overhangs and inactivated by the heat. Then, the overlapping ends anneal, and the DNA polymerase fills the gaps, and the DNA ligase repairs the nicks. Slightly different from these methods, Golden Gate assembly uses type IIS restriction enzymes. Since the type IIS restriction enzymes cleave DNA sequence distant from recognition sequences ^16^, it is possible to leave short single-stranded DNA overhangs of any sequence at the terminal after the cleavage. The DNA fragments with short overhangs are then ligated by T4 DNA ligase. In these methods, after assembly *in vitro,* the assembled plasmids are generally introduced into *E. coli* cells to purify and amplify the desired plasmids. These methods are used for construction of relatively small plasmids, probably due to the low efficiency of introducing large DNA into *E. coli* cells.

On the other hand, methods for assembling DNA inside cells have also been developed. By utilizing the natural transformation ability of *Bacillus subtilis,* a method to construct large DNA has been developed, in which DNA fragments were added stepwise to the genome of *B. subtilis* ^17–19^. *B. subtilis* is not as widespread in many laboratories as *E. coli*, so it has not yet been accessible to many researchers. A method using *Saccharomyces cerevisiae* in constructing large DNA is also attracting attention ^20^, because it was used to build the whole genome of *Mycoplasma genitalium* and *Mycoplasma mycoides* ^21–23^. In this method, multiple DNA fragments are simultaneously introduced into yeast cells and assembled by homologous recombination *in vivo*. The DNA assembly by *S. cerevisiae* was also used for construction of phage genomes for phage therapy ^24,25^. However, the growth rate of *S. cerevisiae* is slower than that of *E. coli,* so it takes several days for colonies to appear, which leads to time loss. For a rapid and simple protocol for introducing multiple DNA fragments into *E. coli* and assembling the DNA fragments, IVA (*<u>i</u>n <u>v</u>ivo* <u>a</u>ssembly) or iVEC (*<u>i</u>n <u>v</u>ivo <u>E</u>. coli* <u>c</u>loning) was also developed ^26–30^. However, it is difficult to assemble large plasmids with this method due to the difficulty in introduction of large or many DNA fragments into *E. coli* cells.

Thus, building large DNA from multiple DNA fragments easily and quickly is still a challenge. This study tackled this challenge by taking advantage of the rapid growth rate of *E. coli* and the ability of bacteriophage to inject long DNA. The sophisticated ability of phages to introduce DNA into their host cells is attractive. This will make it easier and more inexpensive to introduce long DNA into the bacterial cells.

## Results and Discussion

### *In vitro* packaging and transduction of temporarily assembled DNA

I aimed to establish a simple method to assemble large DNA. Therefore, I attempted to combine simple seamless DNA assembly and long DNA introduction by bacteriophage. (Figure 1).

**Figure 1.**
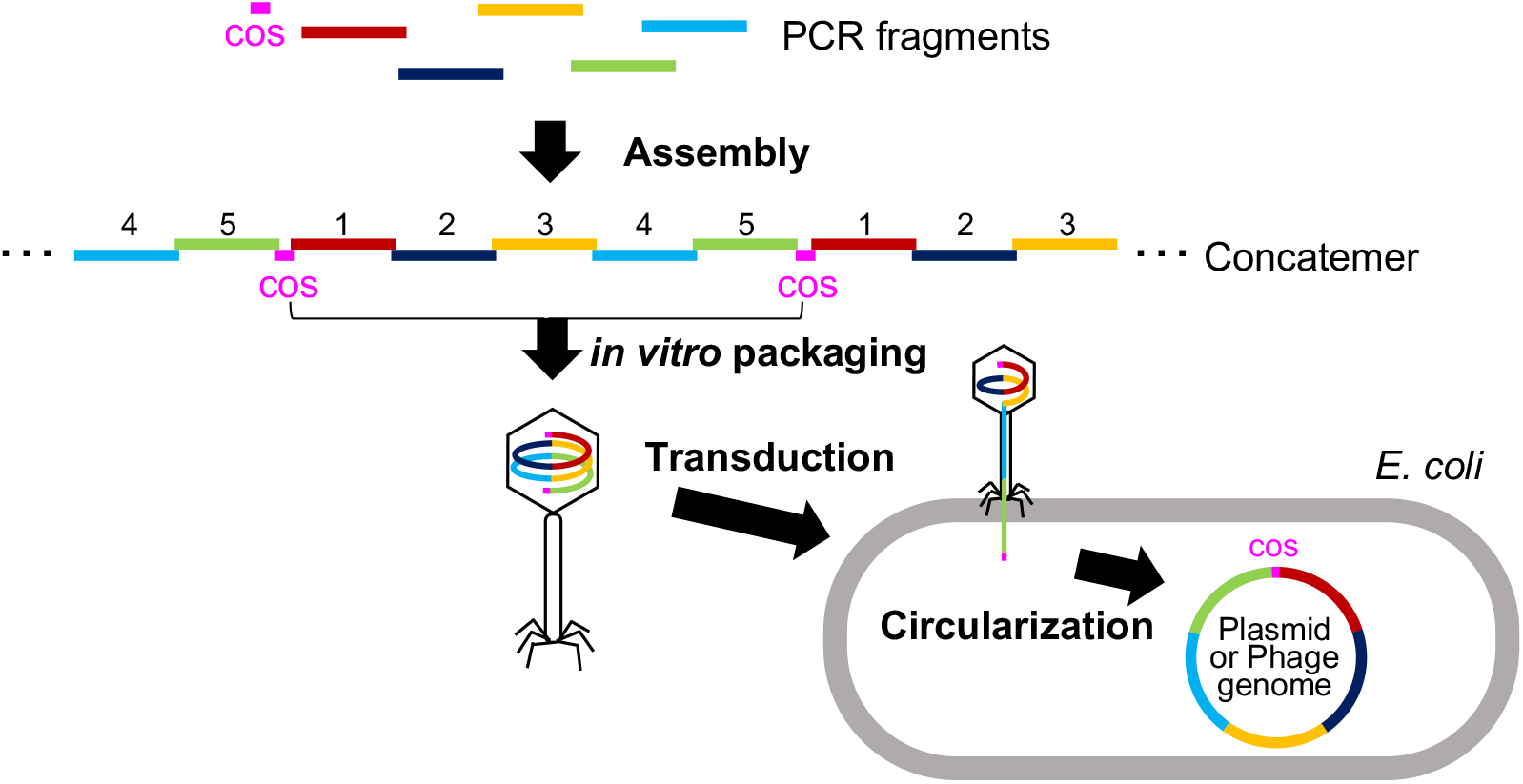
Scheme of iPac (*<u>i</u>n vitro* <u>Pac</u>kaging-assisted DNA assembly). PCR fragments with homologous overlapping ends including the fragments with cos site are assembled by exonuclease treatment and subsequent annealing to form concatemers. The concatemers are packaged into phage particles by *in vitro* packaging. The phage particles are then mixed with *E. coli* cells and the DNA are transduced into cells. The transduced DNA is circularized at the cos site.

To verify whether the temporarily assembled DNA can be introduced into *E. coli* cells using the *in vitro* packaging system of λ phage, I first considered a simple DNA assembly method. PCR fragments can be assembled by using exonuclease III (Exo III) ^31,32^. Exo III has also been shown to contribute to *in vivo* DNA assembly in *E. coli* ^26^. Therefore, a simple assembly of PCR fragments was expected to be performed using Exo III. I found that DNA assembly was possible in just a few minutes by adding DNA fragments that overlaps 50 bp each with adjacent fragments to the reaction solution containing excess Exo III, immediately inactivating the exonuclease at 75°C and returning the DNA fragments to room temperature for annealing (Figure S1). The length of the single-stranded DNA exposed by Exo III treatment cannot be completely controlled. Therefore, it is expected that gaps, flaps or nicks will occur at the junctions of the DNA fragments after Exo III assembly.

### Construction of λ phage genome

The *in vitro* packaging system of λ phage has long been used to introduce DNA into *E. coli* cells ^5,9^. It was reported that the λ phage system allows packaging of heterogeneous DNA with an aberrant structure ^33^. Therefore, I thought it may be possible to package even DNA containing gaps, flaps or nicks. Then, I examined whether it is possible to package the temporarily assembled DNA that was simply annealed after chewing back with Exo III, and to transduce it into *E. coli*.

For this purpose, I attempted to construct a λ phage genome from five PCR fragments of ~9.7 kb (λ_1 to λ_5), each end of which overlaps with the adjacent fragment by 50 bp (Figure 2A, B). The cos site, which is the packaging site of λ phage, is included in λ_1 fragment. The PCR fragments were designed to be circular when assembled so that concatemers as packaging substrates could be formed. It is expected that plaques of λ phage will be detected, only when all these fragments are assembled, packaged into phage capsid, and introduced into *E. coli* cells. No plaque appeared when the PCR fragments without Exo III treatment as a negative control were subjected to a packaging reaction and subsequently mixed with *E. coli* culture (Figure 2C). On the other hand, indeed, many plaques appeared with Exo III treatment (Figure 2D). Approximately 3,000 plaques appeared using at total of 8.6 ng PCR fragments with 5 minutes of heat treatment after Exo III addition and 60 minutes of packaging time. This corresponds to 1 × 10^5^ PFU/μg DNA.

**Figure 2.**
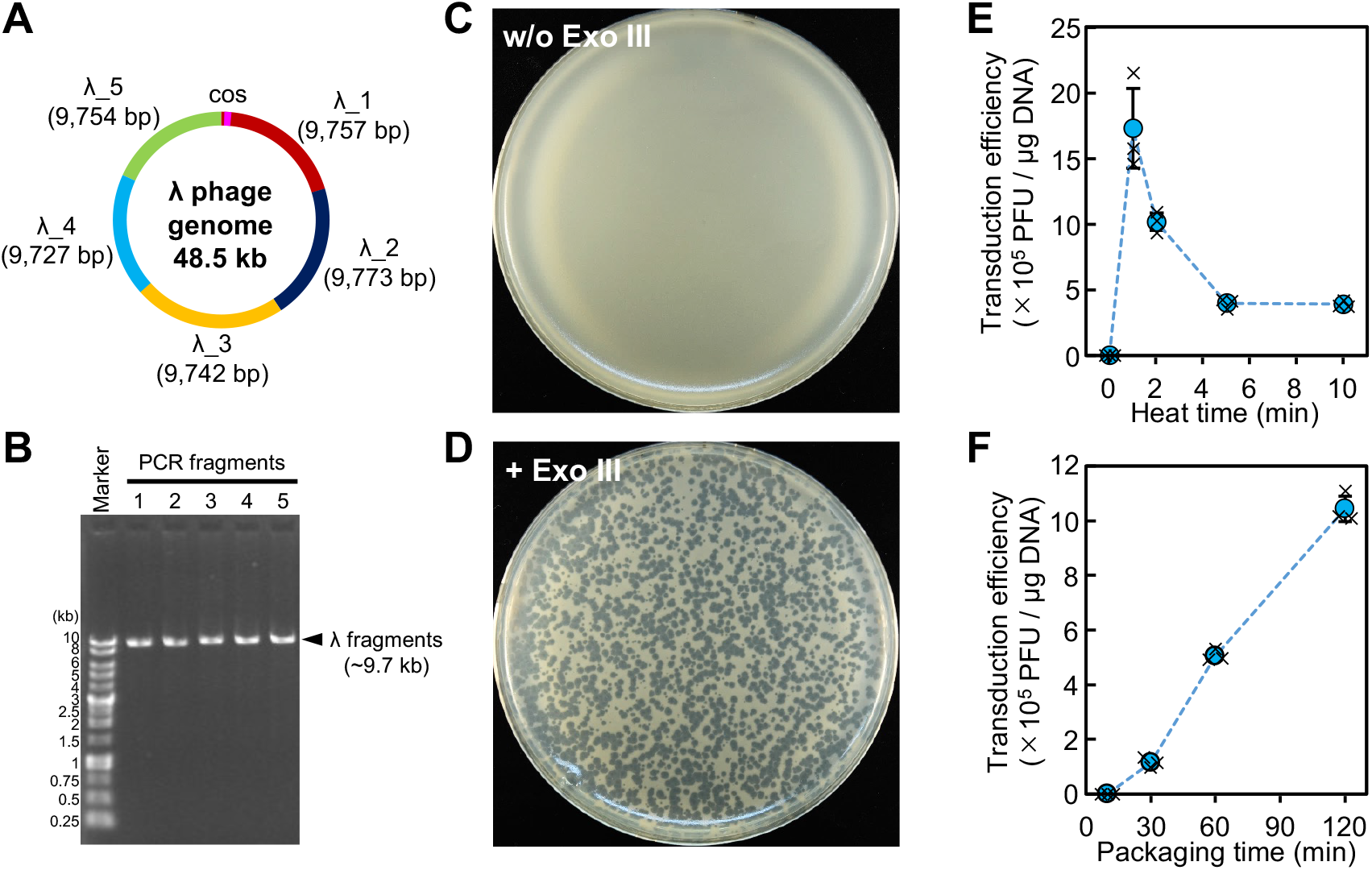
Construction of λ phage by iPac. A. Design of λ phage genome fragments. λ phage genome was divided into 5 fragments of about 9.8 kb (λ_1 to λ_5). The ends of each fragment overlap by 50 bp. The cos site is included in the λ_1 fragment. B. The five PCR-amplified fragments were analyzed by agarose gel electrophoresis. C. D. Agar plates containing indicator *E. coli* cells after *in vitro* packaging and transduction of the DNA fragments without (C) or with (D) Exo III assembly. E. Transduction efficiency according to the heat time after exonuclease treatment. F Transduction efficiency according to the time of *in vitro* packaging. The average plaque forming units of three independent experiments are shown. Error bars and crosses indicate the standard deviations and individual values, respectively.

The number of plaques is expected to increase with improved efficiency in assembly or packaging, so that the plaque number can be a good indicator for better conditions. From this point of view, the most efficient heat time at 75°C for inactivation during Exo III treatment was 1 minute and increasing the heat time reduced the efficiency (Figure 2E). As for the packaging time before mixing with *E. coli* culture, there was almost no plaques in 10 minutes. Enough plaques of about 5 × 10^5^ PFU/μg DNA were appeared with packaging time of 60 minutes, and the efficiency was further improved to 1 × 10^6^ PFU/μg DNA by extending it to 120 minutes (Figure 2F). These results indicate that the temporarily assembled PCR fragments were efficiently packaged by λ phage packaging system and introduced into *E. coli* cells. Here, I call the DNA assembly method using *in vitro* packaging of bacteriophage as iPac (*<u>i</u>n vitro* <u>Pac</u>kaging-assisted DNA assembly). The assembly procedure before *in vitro* packaging will not be essentially limited to the method using Exo III. Various other assembly methods are also expected to work in iPac.

### Modification of λ phage genome by iPac

Next, I examined whether λ phage with various deletion mutants could be easily constructed using iPac by changing the PCR fragments used. The PCR fragments were designed so that the ends of adjacent fragments overlap by 50 bp to delete the target regions (Figure 3A). The deletions were introduced into regions of λ phage genome that are not essential for the lytic cycle of λ phage. As a result of NGS analysis of obtained phage genomes, it was confirmed that the regions of *ea47* (1,690 bp), *ea31-ea59* (2,938 bp), *p35-orf61* (1,843 bp), *orf61-gam* (2,278 bp), *exo* (1,063 bp), *bet* (750 bp), *kil-sieB* (1,754 bp), *rexB-cI* (2,359 bp), *ren-ninI* (3,496 bp) and *bor-p79* (1,877 bp) genes were successfully deleted (Figure 3B). Furthermore, by combining two (*ren-ninI* and *bor-p79*) and four (*kil-sieB, rexB-cI, ren-ninI* and *bor-p79*) of these deletions, I succeeded in construction of λ phage with deletions of total 5,373 bp and 9,486 bp, respectively, by iPac (Figure 3B). In the genome-reduced λ phage, the size of the right arm was reduced from 15.7 kb to 6.3 kb, and the entire genome size was reduced to 39 kb.

**Figure 3.**
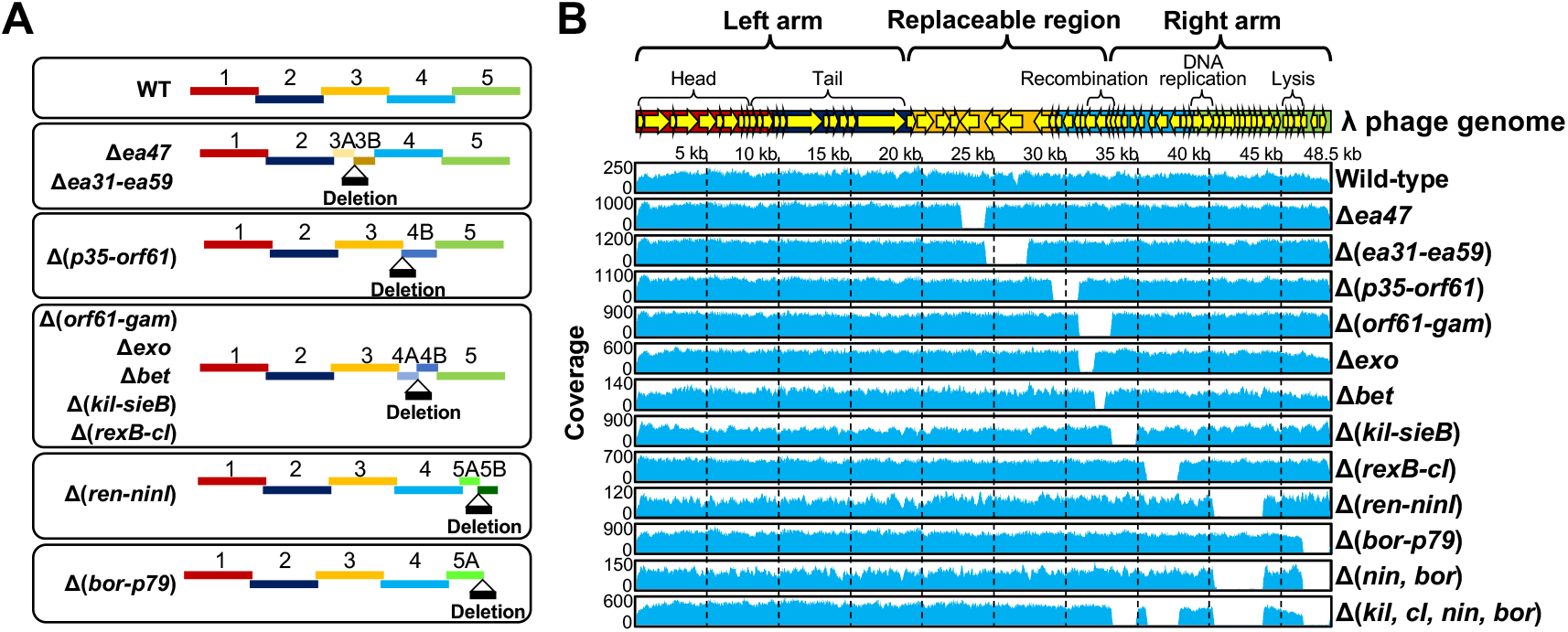
Construction of various deletion mutants of λ phage. A. Design of fragments for deletion construction. Deletions were introduced between the indicated fragments. The ends of each fragment overlap by 50 bp. B. NGS analysis of the constructed λ phage genomes. The structure of the λ phage genome is shown at the top. Reads obtained from NGS analysis of the indicated deletion mutants were mapped to the reference λ phage genome sequence (Genbank accession number: NC_001416).

In the cloning using λ phage vector, the total length of the left and right arms and the DNA fragment to be cloned should be within 52 kb which can be packaged in the capsid of λ phage, so the shorter the left and right arms of λ phage, the longer DNA fragment can be cloned. Therefore, this genome-reduced λ phage will be used as a λ phage vector that is able to clone up to 27 kb, which is longer than the common λEMBL vector that can be cloned up to 23 kb ^34^.

### Construction of various phages by iPac

So far, the λ phage genomes have been generated by using the packaging system of λ phage. Then, can the genomes of other phages be constructed by the λ phage system? To address this question, I selected the four other phages, T1, T3, T7 and φ80 to construct. The genome size of these phages is 39 – 49 kb, which is packageable size with the λ phage system. I designed the genomes of T1/φ80 or T3/T7 to be constructed from 5 or 4 DNA fragments of about 9 - 10 kb, respectively, in addition to a 0.3 kb fragment containing the cos site (Figure 4A-D). The DNA fragments were prepared by PCR (figure 4E-H). The cos site for packaging with λ phage system was placed avoiding regions that might affect transcription; intergenic region between T1p26 (putative tale fiber) and T1p27 (hypothetical 1rich region between T7 promoter A1 and A2 and between φ80 gp35 (hyphothetical protein) and damL (pseudogene) for T1, T3, T7 and φ80, respectively (Figure S2A - D). As a result of introduction of the phage genomes assembled from these DNA fragments into *E. coli* cells by iPac, plaques were obtained in all cases (Figure 4I). The PCR products used for packaging and introduced into *E. coli* cells were 78, 61, 64 and 74 ng for phage T1, T3, T7 and φ80, respectively. And the efficiency was 9,600 ± 1,200, 18,600 ± 1,600, 13,200 ± 1,400 and 829,000 ± 10,500 PFU/μg DNA for phage T1, T3, T7 and φ80, respectively (Table 1).

**Figure 4.**
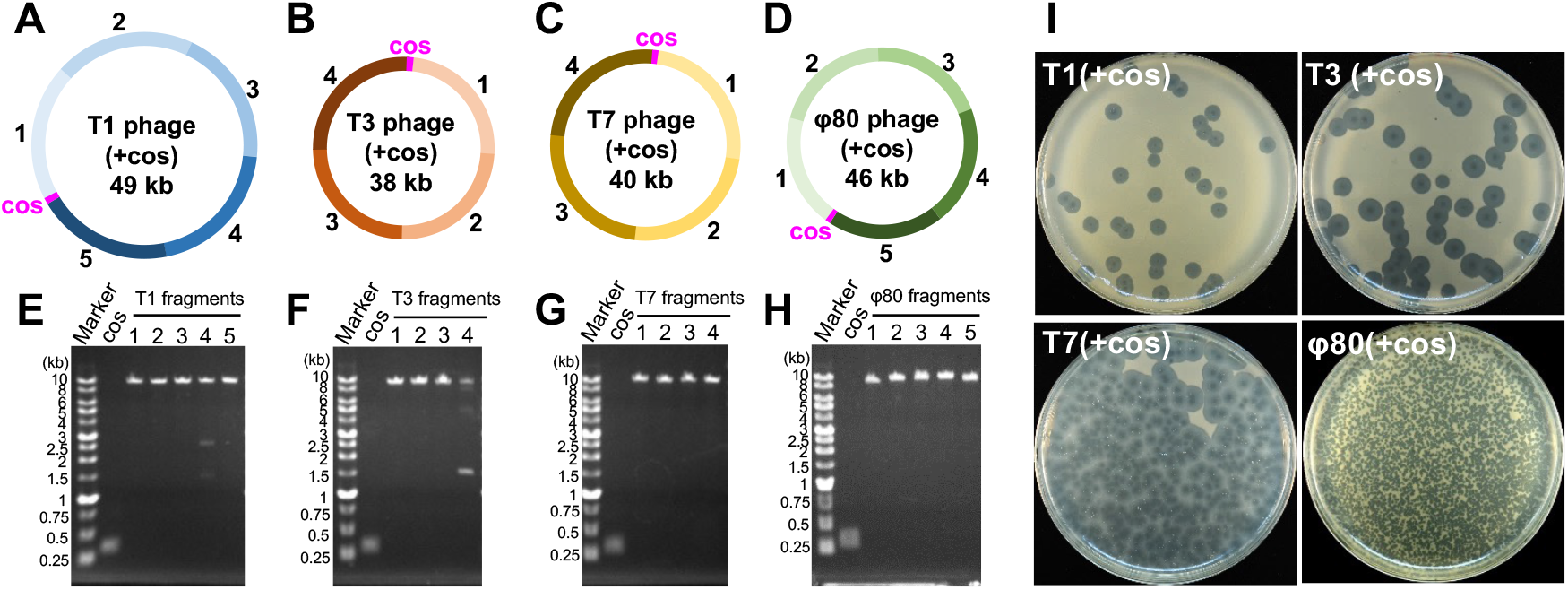
Construction of various phages. A-D. Design of fragments for construction of phage T1(+cos) (A), T3(+cos) (B), T7(+cos) (C) and φ80(+cos) (D) phage genomes. The inserted cos sites are indicated as magenta. E - H. PCR amplified fragments of phage T1 (E), T3 (F), T7 (G) and φ80 (H) genomes were analyzed by agarose gel electrophoresis. I. Plaques of the indicated phages that appeared after the transduction into *E. coli.*

**Table 1:** Number of plaques obtained by iPac in the construction of various phages

| Constructed phage | Number of plaques | | | Average plaque number ± SD | DNA used (ng) | Efficiency (PFU/μg DNA ± SD) |
| --- | --- | --- | --- | --- | --- | --- |
|  | #1 | #2 | #3 |  |  |  |
| T1(+cos) | 33 | 35 | 44 | $37 \pm 5$ | 78 | $9,600 \pm 1,200$ |
| T3(+cos) | 64 | 53 | 54 | $57 \pm 5$ | 61 | $18,600 \pm 1,600$ |
| T7(+cos) | 426 | 415 | 420 | $420 \pm 4$ | 64 | $13,200 \pm 1,400$ |
| ϕ80(+cos) | 3,484 | 2,544 | 3,116 | $3,048 \pm 387$ | 74 | $829,000 \pm 10,500$ |

The phage genomes were recovered from the obtained plaques and analyzed by NGS. For the phage T3, T7 and φ80, the genome was successfully constructed as designed (Figure S2B-D). In the T1 phage, the reads of the inserted cos site were reduced to one third compared to the other genomic regions (Figure S2A). It is considered that, due to the PCR primer design in T1 phage construction, a homologous sequence of 10 bp was generated on both sides of cos site, where deletion of the cos site by homologous recombination occured (Figure S2E). However, the other region of T1 phage genome was successfully constructed. Thus, it is possible to construct the phages other than λ phage by iPac using the packaging system of λ phage.

Thus, iPac has also succeeded in constructing and re-booting the genomes of other phages including lytic phages such as T1, T3 and T7. Because the genome engineering of phages by iPac is simple and rapid, it will be one of the useful options in addition to the conventional phage engineering tools such as for phage therapy ^35^,^36^.

### Construction of a Plasmid by iPac

Next, I examined whether it is also possible to construct a plasmid by iPac. Therefore, I designed a 48 kb plasmid consisting of 4 fragments from P1 phage genome, *cat* (chloramphenicol acetyltransferase) gene and a vector fragment containing ampicillin resistance gene and cos site (Figure 5A). Each fragment was prepared by PCR and was confirmed by agarose gel electrophoresis (Figure 5B). When these PCR fragments were introduced into *E. coli* DH5α strain by iPac and selected only by the ampicillin, more than 1,000 ampicillin-resistant colonies appeared (Figure 5C). When I recovered the plasmids from 20 randomly selected colonies, surprisingly, all 20 recovered plasmids were of the desired size (Figure 5D). Furthermore, as a result of confirming the plasmid structure by restriction fragment length analysis, all the plasmids were correctly constructed (Figure 5E). The average number of colonies in the three independent experiments using 38.2 ng of packaged DNA was 1,785. And therefore, the average transformation efficiency was 4.67 ± 0.85 × 10^4^ CFU/μg DNA (Table 2). Thus, iPac could accurately and efficiently construct large plasmid from multiple PCR fragments.

**Figure 5.**
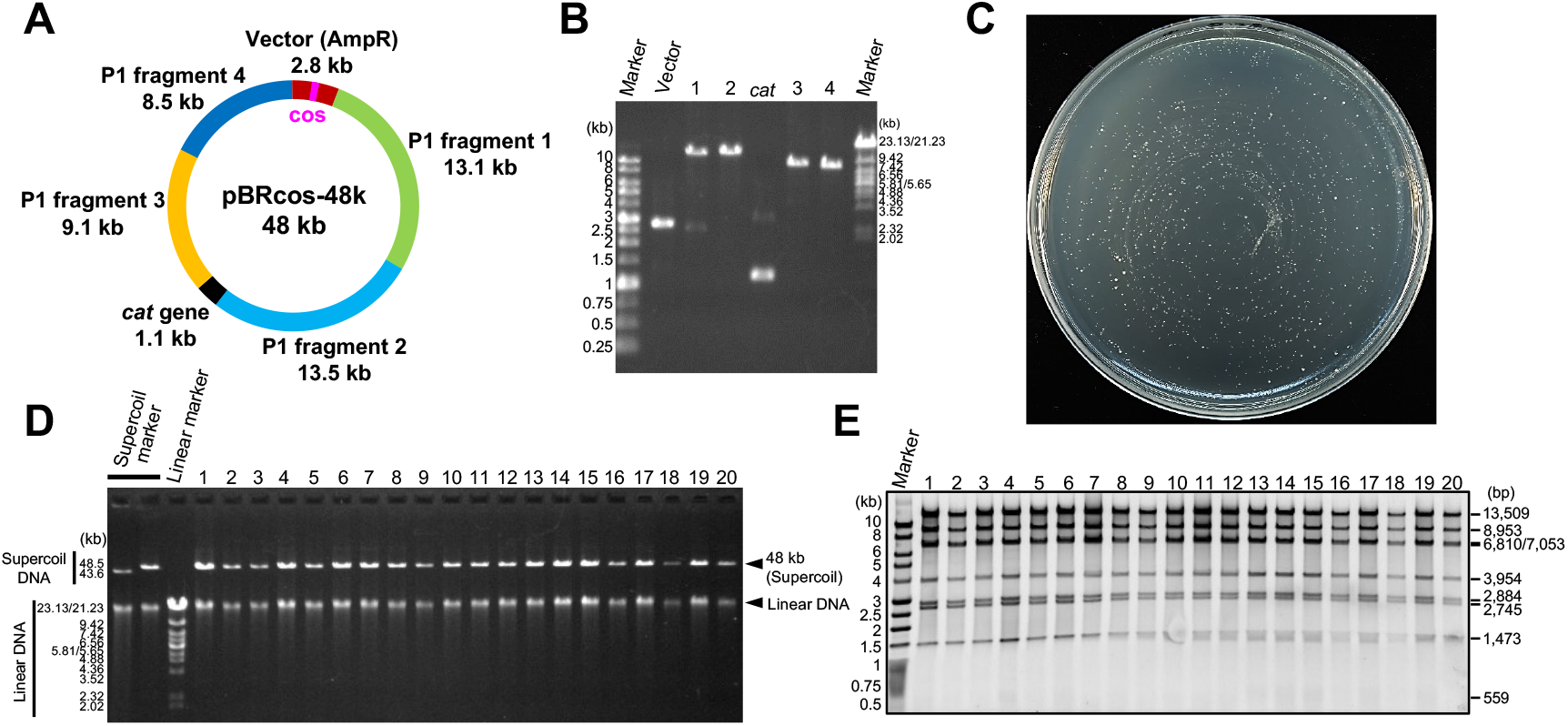
Construction of 48 kb plasmid. A. Design of 48 kb plasmid. pBR322-cos vector, 4 fragments from P1 phage genome and *cat* (chloramphenicol acetyltransferase) gene were used. The ends of adjacent fragments overlap by 50 bp. B. Verification of PCR-amplified DNA fragments by agarose gel electrophoresis. C. Ampicillin-resistant colonies that appeared after the transduction into *E. coli* DH5α strain. D. Agarose gel analysis of the plasmids recovered from 20 randomly selected colonies. Linear DNA is due to contamination of genomic DNA or double-strand breaks of the plasmid DNA. E. Verification of plasmid construction by a restriction enzyme. The plasmids digested by Nde I were analyzed by agarose gel electrophoresis. The expected sizes of the restriction fragments are indicated on the right.

**Table 2:** Number of colonies obtained by iPac in the construction of 48 kb plasmid

| Number of colonies | | | Average colony<br>number ± SD | DNA used<br>(ng) | Efficiency<br>(CFU/μg DNA) |
| --- | --- | --- | --- | --- | --- |
| #1 | #2 | #3 |  |  |  |
| 1,821 | 2,161 | 1,372 | 1,785 ± 323 | 38.2 | 46,700 ± 8,500 |

The problem with the current iPac system is that the DNA size that can be packaged in the capsid of λ phage is limited to about 38 – 52 kb. By using the *in vitro* packaging system of phages with larger genomes, such as P1 or T4 phage with the genome size of 94 and 169 kb, respectively, it will be possible to construct DNA larger than the packaging limit of λ phage system. Furthermore, by using packaging system of phages that infect other bacteria, iPac system may be applicable to other bacteria. On the contrary, limiting the size of the packaged DNA can also be beneficial. Generally, in the assembly of multiple DNA fragments, it is inevitable that the wrong assembly results in the construction of the wrong size plasmid. In iPac system, the assembled DNA that is too small or too large to complete packaging into the phage capsid is excluded during the packaging process. It is considered that this size elimination mechanism made it possible to construct the plasmid with extremely high accuracy (Figure 5D, 5E). In this study, up to 6 DNA fragments were assembled by iPac. There is room for further research as to whether it can be applied to the assembly of more fragments. By Golden Gate assembly, 40 kb genome of T7 phage was constructed from 52 DNA fragments ^37^. Combining iPac with Golden Gate assembly may improve the efficiency of multifragment construction.

When constructing a large plasmid from multiple DNA fragments, the concentration of the DNA fragments should be lowered to increase the efficiency of self-circularization by joining intramolecular ends, although the efficiency of assembly between each DNA fragment decreases. On the other hand, the efficiency of the assembly rises by increasing the concentration of the DNA fragments. But in this case, the concatemers produced by intermolecular end joining are more likely to form instead of circular plasmids. In iPac system, the concatemer is the substrate for packaging, and the circularization occurs inside the cells after transduction. Therefore, this conflicting problem is solved, enabling highly efficient assembly. In addition, iPac doesn’t require special equipment such as electroporator or high-performance competent cells, which are generally required for the introduction of large DNA into *E. coli*. The transduction in iPac is completed simply by mixing the *E. coli* cells prepared from the overnight culture and is performed at low cost.

To use the iPac system presented here, the packaging extract and the cos sequence is required. There are commercially available *in vitro* packaging kits, but the packaging extract can also be homemade ^6,38,39^. The homemade packaging extract will further reduce the cost for iPac. Recently, an *in vitro* transcription / translation system have been put into practical use for phage reconstitution ^40^, which may allow the reconstitution of the packaging extracts not limited to λ phage as well. The cos sequence is required for the packaging using the λ phage system. The cos site is included in some of the commonly used vectors such as BAC vector, pBe1oBAC11 and fosmid vector, pFOS1 ^41^. Therefore, plasmid construction by iPac can be readily performed using these vectors without preparing the cos fragment individually.

## Conclusions

I have presented that it is possible to construct large DNA rapidly and accurately using only the materials accessible to many researchers such as *E. coli* and *in vitro* packaging system. This method will lower the barriers to entry for research using larger DNA.

## Methods

### Bacterial strains and medium

*Escherichia coli* strains used for this study were listed on Table 3. SN1171, BL21(DE3) and DH5α were used for the recipient strains for transduction. LB broth (1% trypton, 0.5% yeast extract, 1% NaCl) (Nacalai Tesque) was used for liquid culture. The agar plates were made by adding 1.5% agar to LB broth. 50 μg/mL of Ampicillin was added to the medium when antibiotic resistance selection is needed. MgSO_4_ with final concentration of 10 mM and 0.7% agar was added to LB broth to make soft agar for plaque assay.

**Table 3:**
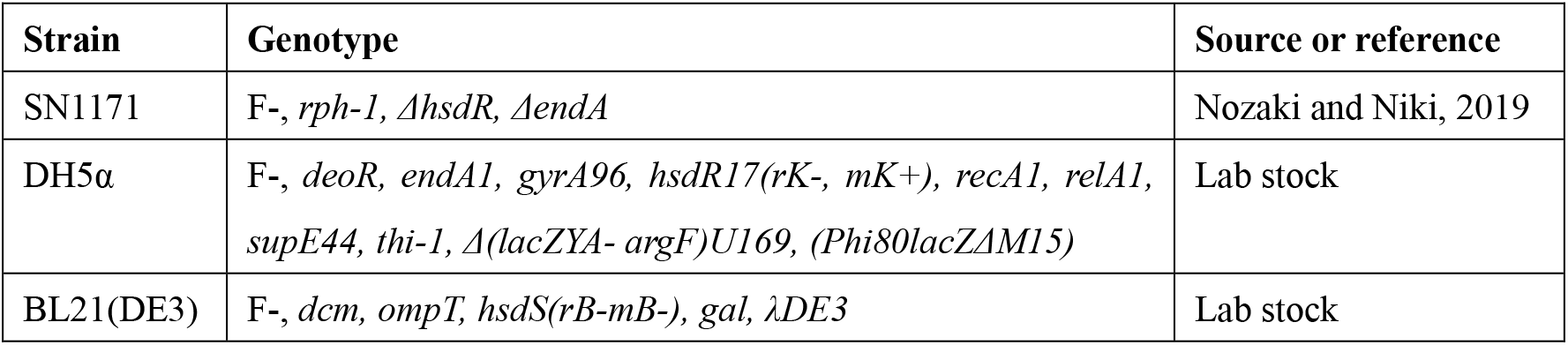
Bacterial strains

### Bacteriophages

Genome of λ phage (*c1857, Sam7*) was purchased from NIPPON GENE. The lambda phage strain (*cI857*) in which *Sam7* mutation was reverted to the wild type was obtained from a plaque that appeared after infection of λ *cI857 Sam7* to *E. coli* SN1171 strain without the amber suppressor mutation. Phages T1, T3, T7, φ80 and P1kc were distributed from Biological Resource Center, NITE, Japan

### Plasmids

pBR322-cos, a plasmid containing the packaging site of λ phage, was constructed by inserting cos sequence between replication origin and β-lactamase gene of pBR322 ^42^ by iVEC method as described previously ^26^. For amplification of chloramphenicol acetyltransferase (*cat*) gene, pKD3 ^43^ was used as a template.

### Preparation of PCR products

PCR was carried out with KOD One DNA polymerase (TOYOBO) according to the manufacturer’s instruction. The oligonucleotide primers were designed so that the DNA fragments to be assembled overlap each other by 50 bp at the ends. The oligonucleotide primer sequences, and combinations of the primers and templates are listed in Table S1 and S2. The PCR products were purified using NucleoSpin PCR clean-up kit (Takara).

### *in vitro* packaging-assisted DNA assembly and transduction

For preparing the assembly reaction, 2x Assembly Solution (20 mM Tris-HCl (pH 7.9), 100 mM of NaCl, 20 mM MgCl_2_, 10% (w/v) PEG8,000, 2 mM Dithiothreitol, 1.2 U/μL Exonuclease III (Takara)) and an equal amount of 4.8 nM each of DNA fragments was mixed to carry out the assembly reaction. In practice, 5 μL volume of the reaction (2.5 μL of 2x Assembly Solution + 2.5 μL of DNA mix) was prepared on ice and incubated at 75°C for 1 minute immediately after the addition of DNA mix. The combinations of DNA fragments to be assembled for each construct were shown in Table S3. Then, the reaction was incubated at 25°C for 5 minutes to allow annealing at exposed single-stranded ends. 0.5 μL of assembly reaction was mixed with 4 μL of packaging extract from *in vitro* packaging kit, LAMBDA INN (NIPPON GENE) and stand for 60 – 120 minutes at 25°C for *in vitro* packaging. For transduction, 1 mL of overnight culture of *E. coli* cells were collected by centrifugation (5,000 g for 1 min). For construction of phages, *E. coli* strain SN1171 was used except for T3 phage construction in which BL21(DE3) was used. For construction of the 48 kb plasmid, DH5α was used. The collected cells were resuspended in 500 μL of 10 mM MgSO4. 100 μL of the cell suspension were mixed with 4.5 μL of the *in vitro* packaging reaction and incubated at 25°C for 15 min. For plaque formation in phage construction, the transduced cells were mixed with 2.5 mL of soft agar prewarmed at 50°C and spread on LB agar plate. For plasmid construction, the transduced cells were spread on LB agar plate with 50 μg/mL ampicillin. The agar plates were incubated at 37°C for overnight and plaques or colonies that emerged on the plates were analyzed.

### Next-Generation Sequencing (NGS) analysis

The libraries from recovered phage genomes for NGS analysis were prepared with Nextera DNA Flex Library Prep kit (illumina). The libraries were sequenced by iSeq 100 system (illumina). The obtained reads were analyzed by Geneious software (Biomatters Ltd.) with the reference genome sequences of phages; λ, T1, T3, T7 and phi80 (Genbank accession numbers NC_001416, NC_005833, NC_003298, NC_001604 and NC_021190, respectively).

### Plasmids analysis

Plasmids constructed by iPac were recovered with Qiaprep spin miniprep kit (Qiagen). The purified plasmids were analyzed by 0.7% agarose gel electrophoresis. The plasmid structure was verified by restriction enzyme treatment. For restriction enzyme digestion, 2.5 uL of purified plasmids were digested with Nde I (New England BioLabs) and analyzed by 0.7% agarose gel electrophoresis.

## Supporting information

Tables S1, S2, S3 and Figures S1, S2, and will be used for the link to the file on the preprint site.

## Abbreviations

PCR: Polymerase Chain Reaction
NGS: Next Generation Sequencing
PEG: Polyethylene Glycol
PFU: Plaque Forming Unit
CFU: Colony Forming Unit

## Author information

Shingo Nozaki

Graduate School of Advanced Science and Engineering, Hiroshima University, 1-4-1 Kagamiyama, Higashi-Hiroshima, Hiroshima 739-8527, Japan

## Acknowledgement

I am grateful to Prof. Masahiro Su’etsugu for his help, including research space and equipment, and Emi Hagiuda for technical assistance in NGS analysis. I would also like to thank Dr. Takahito Mukai and Dr. Shota Suzuki for valuable discussions. For the distribution of the *E. coli* strain and the phages, I would like to thank NBRP *E. coli* at NIG and NBRC at NITE, respectively. This work was supported by JST PRESTO Grant Number JPMJPR19K5 and JSPS KAKENHI Grant Number JP19K05782.

## Notes

### Competing Interest Statement

The authors have declared no competing interest.

## References

(1) Richardson, S. M.; Mitchell, L. A.; Stracquadanio, G.; Yang, K.; Dymond, J. S.; DiCarlo, J. E.; Lee, D.; Huang, C. L. V.; Chandrasegaran, S.; Cai, Y.; Boeke, J. D.; Bader, J. S. Design of a Synthetic Yeast Genome. Science 2017, 355 (6329), 1040–1044. https://doi.org/10.1126/science.aaf4557.

(2) Fredens, J.; Wang, K.; de la Torre, D.; Funke, L. F. H.; Robertson, W. E.; Christova, Y.; Chia, T.; Schmied, W. H.; Dunkelmann, D. L.; Beránek, V.; Uttamapinant, C.; Llamazares, A. G.; Elliott, T. S.; Chin, J. W. Total Synthesis of *Escherichia Coli* with a Recoded Genome. Nature 2019, 569 (7757), 514–518. https://doi.org/10.1038/s41586-019-1192-5.

(3) Cohen, S. N.; Chang, A. C. Y.; Boyer, H. W.; Helling, R. B. Construction of Biologically Functional Bacterial Plasmids *In Vitro*. Proceedings of the National Academy of Sciences 1973, 70 (11), 3240–3244. https://doi.org/10.1073/pnas.70.11.3240.

(4) Murray, N. E.; Murray, K. Manipulation of Restriction Targets in Phage λ to Form Receptor Chromosomes for DNA Fragments. Nature 1974, 251 (5475), 476–481. https://doi.org/10.1038/251476a0.

(5) Hohn, B.; Wurtz, M.; Klein, B.; Lustig, A.; Hohn, T. Phage Lambda DNA Packaging, *in Vitro*. Journal of Supramolecular Structure 1974, 2 (2–4), 302–317. https://doi.org/10.1002/jss.400020220.

(6) Hohn, B.; Murray, K. Packaging Recombinant DNA Molecules into Bacteriophage Particles *in Vitro*. Proceedings of the National Academy of Sciences 1977, 74 (8), 3259–3263. https://doi.org/10.1073/pnas.74.8.3259.

(7) Sternberg, N.; Tiemeier, D.; Enquist, L. In Vitro Packaging of a λ Dam Vector Containing EcoRI DNA Fragments of *Escherichia Coli* and Phage P1. Gene 1977, 1 (3–4), 255–280. https://doi.org/10.1016/0378-1119(77)90049-X.

(8) Thomas, M.; Cameron, J. R.; Davis, R. W. Viable Molecular Hybrids of Bacteriophage Lambda and Eukaryotic DNA. Proceedings of the National Academy of Sciences 1974, 71 (11), 4579–4583. https://doi.org/10.1073/pnas.71.11.4579.

(9) Collins, J.; Hohn, B. Cosmids: A Type of Plasmid Gene-Cloning Vector That Is Packageable *in Vitro* in Bacteriophage Lambda Heads. Proceedings of the National Academy of Sciences 1978, 75 (9), 4242–4246. https://doi.org/10.1073/pnas.75.9.4242.

(10) Tao, W.; Chen, L.; Zhao, C.; Wu, J.; Yan, D.; Deng, Z.; Sun, Y. *In Vitro* Packaging Mediated One-Step Targeted Cloning of Natural Product Pathway. ACS Synthetic Biology 2019, 8 (9), 1991–1997. https://doi.org/10.1021/acssynbio.9b00248.

(11) Li, M. Z.; Elledge, S. J. Harnessing Homologous Recombination *in Vitro* to Generate Recombinant DNA via SLIC. Nature Methods 2007, 4 (3), 251–256. https://doi.org/10.1038/nmeth1010.

(12) Zhu, B.; Cai, G.; Hall, E. O.; Freeman, G. J. In-Fusion^TM^ Assembly: Seamless Engineering of Multidomain Fusion Proteins, Modular Vectors, and Mutations. Biotechniques 2007, 43 (3), 354–359. https://doi.org/10.2144/000112536.

(13) Engler, C.; Gruetzner, R.; Kandzia, R.; Marillonnet, S. Golden Gate Shuffling: A One-Pot DNA Shuffling Method Based on Type IIs Restriction Enzymes. PLoS ONE 2009, 4 (5), e5553. https://doi.org/10.1371/journal.pone.0005553.

(14) Gibson, D. G.; Young, L.; Chuang, R. Y.; Venter, J. C.; Hutchison, C. A.; Smith, H. O. Enzymatic Assembly of DNA Molecules up to Several Hundred Kilobases. Nature Methods 2009, 6 (5), 343–345. https://doi.org/10.1038/nmeth.1318.

(15) Gibson, D. G. Enzymatic Assembly of Overlapping DNA Fragments. In Methods in Enzymology; Academic Press Inc., 2011; Vol. 498, pp 349–361. https://doi.org/10.1016/B978-0-12-385120-8.00015-2.

(16) Szybalski, W.; Kim, S. C.; Hasan, N.; Podhajska, A. J. Class-IIS Restriction Enzymes — a Review. Gene 1991, 100, 13–26. https://doi.org/10.1016/0378-1119(91)90345-C.

(17) Tsuge, K.; Matsui, K.; Itaya, M. One Step Assembly of Multiple DNA Fragments with a Designed Order and Orientation in Bacillus Subtilis Plasmid. Nucleic Acids Res 2003, 31 (21). https://doi.org/10.1093/nar/gng133.

(18) Itaya, M.; Tsuge, K.; Koizumi, M.; Fujita, K. Combining Two Genomes in One Cell: Stable Cloning of the Synechocystis PCC6803 Genome in the Bacillus Subtilis 168 Genome. Proceedings of the National Academy of Sciences 2005, 102 (44), 15971–15976. https://doi.org/10.1073/pnas.0503868102.

(19) Itaya, M.; Fujita, K.; Kuroki, A.; Tsuge, K. Bottom-up Genome Assembly Using the Bacillus Subtilis Genome Vector. Nature Methods 2008, 5 (1), 41–43. https://doi.org/10.1038/nmeth1143.

(20) Kouprina, N.; Larionov, V. Transformation-Associated Recombination (TAR) Cloning for Genomics Studies and Synthetic Biology. Chromosoma 2016, 125 (4), 621–632. https://doi.org/10.1007/s00412-016-0588-3.

(21) Gibson, D. G.; Glass, J. I.; Lartigue, C.; Noskov, V. N.; Chuang, R.-Y.; Algire, M. A.; Benders, G. A.; Montague, M. G.; Ma, L.; Moodie, M. M.; Merryman, C.; Vashee, S.; Krishnakumar, R.; Assad-Garcia, N.; Andrews-Pfannkoch, C.; Denisova, E. A.; Young, L.; Qi, Z.-Q.; Segall-Shapiro, T. H.; Calvey, C. H.; Parmar, P. P.; Hutchison, C. A.; Smith, H. O.; Venter, J. C. Creation of a Bacterial Cell Controlled by a Chemically Synthesized Genome. Science (1979) 2010, 329 (5987), 52–56. https://doi.org/10.1126/science.1190719.

(22) Gibson, D. G.; Benders, G. A.; Andrews-Pfannkoch, C.; Denisova, E. A.; Baden-Tillson, H.; Zaveri, J.; Stockwell, T. B.; Brownley, A.; Thomas, D. W.; Algire, M. A.; Merryman, C.; Young, L.; Noskov, V. N.; Glass, J. I.; Venter, J. C.; Hutchison, C. A.; Smith, H. O. Complete Chemical Synthesis, Assembly, and Cloning of a *Mycoplasma Genitalium* Genome. Science (1979) 2008, 319 (5867), 1215–1220. https://doi.org/10.1126/science.1151721.

(23) Gibson, D. G.; Benders, G. A.; Axelrod, K. C.; Zaveri, J.; Algire, M. A.; Moodie, M.; Montague, M. G.; Venter, J. C.; Smith, H. O.; Hutchison, C. A. One-Step Assembly in Yeast of 25 Overlapping DNA Fragments to Form a Complete Synthetic Mycoplasma Genitalium Genome. Proceedings of the National Academy of Sciences 2008, 105 (51), 20404–20409. https://doi.org/10.1073/pnas.0811011106.

(24) Ando, H.; Lemire, S.; Pires, D. P.; Lu, T. K. Engineering Modular Viral Scaffolds for Targeted Bacterial Population Editing. Cell Systems 2015, 1 (3), 187–196. https://doi.org/10.1016/j.cels.2015.08.013.

(25) Pires, D. P.; Monteiro, R.; Mil-Homens, D.; Fialho, A.; Lu, T. K.; Azeredo, J. Designing P. Aeruginosa Synthetic Phages with Reduced Genomes. Scientific Reports 2021, 11 (1), 2164. https://doi.org/10.1038/s41598-021-81580-2.

(26) Nozaki, S.; Niki, H. Exonuclease III (XthA) Enforces *In Vivo* DNA Cloning of *Escherichia Coli* To Create Cohesive Ends. Journal of Bacteriology 2019, 201 (5). https://doi.org/10.1128/JB.00660-18.

(27) García-Nafría, J.; Watson, J. F.; Greger, I. H. IVA Cloning: A Single-Tube Universal Cloning System Exploiting Bacterial In Vivo Assembly. Scientific Reports 2016, 6 (June), 1–12. https://doi.org/10.1038/srep27459.

(28) Kostylev, M.; Otwell, A. E.; Richardson, R. E.; Suzuki, Y. Cloning Should Be Simple: *Escherichia Coli* DH5á-Mediated Assembly of Multiple DNA Fragments with Short End Homologies. PLoS ONE 2015, 10 (9), 1–15. https://doi.org/10.1371/journal.pone.0137466.

(29) Jacobus, A. P.; Gross, J. Optimal Cloning of PCR Fragments by Homologous Recombination in *Escherichia Coli*. PLoS ONE 2015, 10 (3), 1–17. https://doi.org/10.1371/journal.pone.0119221.

(30) Watson, J. F.; García-Nafría, J. In Vivo DNA Assembly Using Common Laboratory Bacteria: A Re-Emerging Tool to Simplify Molecular Cloning. Journal of Biological Chemistry 2019, 294 (42), 15271–15281. https://doi.org/10.1074/jbc.REV119.009109.

(31) Hsiao, K.-C. Exonuclease III Induced Ligase-Free Directional Subcloning of PCR Products; 1993; Vol. 21.

(32) Kaluz, S.; Kölble, K.; Reid, K. B. M. Directional Cloning of PCR Products Using Exonuclease III. Nucleic Acids Research 1992, 20 (16), 4369–4370. https://doi.org/10.1093/nar/20.16.4369.

(33) Pearson, R. K.; Fox, M. S. Effects of DNA Heterologies on Bacteriophage Lambda Packaging. Genetics 1988, 118 (1), 5–12. https://doi.org/10.1093/genetics/118.1.5.

(34) Frischauf, A.-M.; Lehrach, H.; Poustka, A.; Murray, N. Lambda Replacement Vectors Carrying Polylinker Sequences. Journal of Molecular Biology 1983, 170 (4), 827–842. https://doi.org/10.1016/S0022-2836(83)80190-9.

(35) Kilcher, S.; Loessner, M. J. Engineering Bacteriophages as Versatile Biologics. Trends in Microbiology 2019, 27 (4), 355–367. https://doi.org/10.1016/j.tim.2018.09.006.

(36) Chen, Y.; Batra, H.; Dong, J.; Chen, C.; Rao, V. B.; Tao, P. Genetic Engineering of Bacteriophages Against Infectious Diseases. Frontiers in Microbiology 2019, 10. https://doi.org/10.3389/fmicb.2019.00954.

(37) Pryor, J. M.; Potapov, V.; Bilotti, K.; Pokhrel, N.; Lohman, G. J. S. Rapid 40 Kb Genome Construction from 52 Parts through Data-Optimized Assembly Design. ACS Synthetic Biology 2022, 11 (6), 2036–2042. https://doi.org/10.1021/acssynbio.1c00525.

(38) Gunther, E. J.; Murray, N. E.; Glazer, P. M. High Efficiency, Restriction-Deficient *in Vitro* Packaging Extracts for Bacteriophage Lambda DNA Using a New *E.Coli* Lysogen. Nucleic Acids Research 1993, 21 (16), 3903–3904. https://doi.org/10.1093/nar/21.16.3903.

(39) Rosenberg, S. M.; Stahl, M. M.; Kobayashi, I.; Stahl, F. W. Improved *in Vitro* Packaging of Coliphage Lambda DNA: A One-Strain System Free from Endogenous Phage. Gene 1985, 38 (1–3), 165–175. https://doi.org/10.1016/0378-1119(85)90215-X.

(40) Rustad, M.; Eastlund, A.; Jardine, P.; Noireaux, V. Cell-Free TXTL Synthesis of Infectious Bacteriophage T4 in a Single Test Tube Reaction. Synthetic Biology 2018, 3 (1). https://doi.org/10.1093/synbio/ysy002.

(41) Shizuya, H.; Birren, B.; Kim, U. J.; Mancino, V.; Slepak, T.; Tachiiri, Y.; Simon, M. Cloning and Stable Maintenance of 300-Kilobase-Pair Fragments of Human DNA in *Escherichia Coli* Using an F-Factor-Based Vector. Proceedings of the National Academy of Sciences 1992, 89 (18), 8794–8797. https://doi.org/10.1073/pnas.89.18.8794.

(42) Bolivar, F.; Rodriguez, R. L.; Greene, P. J.; Betlach, M. C.; Heyneker, H. L.; Boyer, H. W.; Crosa, J. H.; Falkow, S. Construction and Characterization of New Cloning Vehicle. II. A Multipurpose Cloning System. Gene 1977, 2 (2), 95–113. https://doi.org/10.1016/0378-1119(77)90000-2.

(43) Datsenko, K. A.; Wanner, B. L. One-Step Inactivation of Chromosomal Genes in *Escherichia Coli* K-12 Using PCR Products. Proceedings of the National Academy of Sciences 2000, 97 (12), 6640–6645. https://doi.org/10.1073/pnas.120163297.

