## Supplementary material for "Rapid and accurate assembly of large DNA assisted by *in vitro* packaging of bacteriophage": Tables S1, S2, S3 and Figures S1, S2, and will be used for the link to the file on the preprint site.

Shingo Nozaki ^1, 2^

^1^Department of Life Science, College of Science, Rikkyo University, Tokyo 171-8501, Japan

^2^Graduate School of Advanced Science and Engineering, Hiroshima University, Hiroshima 739-8527, Japan

**Table S1: Oligonucleotide primers used for PCR**

| **Name** | **Oligonucleotide sequence (5' - 3')** |
| --- | --- |
| ONR454 | CCACGCACGTTGTGATATGT |
| ONR178 | GACGGTAATTTCTGCAACC |
| ONR179 | CGGTTGTATCCGGTAATGGTGAG |
| ONR180 | ATACCCGGGAGTGATTTCC |
| ONR709 | GGTCCGGCAGTACAATGGATT |
| ONR182 | TAATTGCGGAGACTTTGCGATG |
| ONR183 | GTGCGTTCCACTCCTGAAG |
| ONR184 | GCGTAACCATCATCGAGATCTG |
| ONR185 | AACTTTGCCGGACAGGAGC |
| ONR455 | GGACCCGTAAAGTGATAATG |
| ONR513 | CAAAATCCGGTAGTAACTTGCTAACTGGCGGTGATGTAAACACTA |
| ONR514 | GTTCATAGTGTTTACATCACCGCCAGTTAGCAAGTTACTACCGGA |
| ONR515 | CCCACACCCAGCATGCATACCTTTCTCCGGTAGTAACTTGCTAAC |
| ONR516 | AATTGGTTAGCAAGTTACTACCGGAGAAAGGTATGCATGCTGGGT |
| ONR519 | CTGTGTGCTTATGCTTGCCGACATAATTGCGGAGACTTTGCGATG |
| ONR520 | AAGTACATCGCAAAGTCTCCGCAATTATGTCGGCAAGCATAAGCACA |
| ONR517 | ATGTTTCGTGAAGCCGTCGACGCTTTGTGCTTATGCTTGCCGACA |
| ONR518 | TCCCATGTCGGCAAGCATAAGCACAAAGCGTCGACGGCTTCACGA |
| ONR522 | CGCAGAGCAGAAGGTGGCAGCATGATGTGCTTATGCTTGCCGACA |
| ONR521 | TCCCATGTCGGCAAGCATAAGCACATCATGCTGCCACCTTCTGCT |
| ONR526 | ATAAAACGAATGAGTACTGCACTCGAGCAGAAGGTGGCAGCATGAC |
| ONR525 | CGGTGTCATGCTGCCACCTTCTGCTCGAGTGCAGTACTCATTCGT |
| ONR645 | TAGTTACTTAGATATTGGCCTTGGCTCGTGAAGCCGTCGACGCTT |
| ONR646 | TTTATAAGCGTCGACGGCTTCACGAGCCAAGGCCAATATCTAAGT |
| ONR647 | GAACTTGGCTTATCCCAGGAATCTGATTGGCCGTAAGTGCGATTC |
| ONR648 | ATCCGGAATCGCACTTACGGCCAATCAGATTCCTGGGATAAGCCA |
| ONR610 | TGGGGACGCATAATAGCTTCTGTGCGCAGTGTTCTGCGGAAACCT |
| ONR611 | ATGTTAGGTTTCCGCAGAACACTGCGCACAGAAGCTATTATGCGT |
| ONR505 | CATTTACGAATGTTTGCTGGGTTTCCCACGCACGTTGTGATATGT |
| ONR506 | CATCTACATATCACAACGTGCGTGGGAAACCCAGCAAACATTCGT |
| ONR748 | TATTACTTCATCTCAATTGCCTTCGCCACGCACGTTGTGATATGT |
| ONR749 | ACGTAGACTCATTTCCGAAGGCAATATGTCAGCCAGCTGCTTTTTG |
| ONR750 | TCAACAAAAAGCAGCTGGCTGACATATTGCCTTCGGAAATGAGTC |
| ONR751 | CTGACCATTCAATCCTCTGC |
| ONR752 | GAGCTTTGACACGTTTGAGG |
| ONR753 | TAATCAAGACCCATGCAGTC |
| ONR754 | GAACTCTTGTCCGCTACGAG |
| ONR755 | CCTGTTTCATATCCTCTTCG |
| ONR756 | GGGTTCAAATGAGACTGGAG |
| ONR757 | GCCGAACTTACTACCATCAAC |
| ONR758 | ATCCGGGAGAGATAATCACTC |
| ONR759 | CATCTACATATCACAACGTGCGTGGCGAAGGCAATTGAGATGAAG |
| ONR760 | TAAAGAGCGACGCTATCTTAAAGACCCACGCACGTTGTGATATGT |
| ONR761 | TAACAGTGGGTCTATCGGTCTGGTTCCATTGTTCATTCCACGGAC |
| ONR762 | TTTTTGTCCGTGGAATGAACAATGGAACCAGACCGATAGACCCAC |
| ONR763 | TTCGCCAAAGTTCATAGTCG |
| ONR764 | AAAGCAGCTTGAGAGTAAGG |
| ONR765 | ATAAGCGTGCTTCTATCTGG |
| ONR766 | TGACTTTACCGTAGCGAAAG |
| ONR767 | TGGTGTCCATGAGGTAAATG |
| ONR768 | GCAACTGACCATGTATTCTC |
| ONR769 | CATCTACATATCACAACGTGCGTGGGTCTTTAAGATAGCGTCGCT |
| ONR770 | CTATAGGATACTTACAGCCATCGAGCCACGCACGTTGTGATATGT |
| ONR771 | TGGGATGGCTATTCGCCGTGTCCCTCCATTGTTCATTCCACGGAC |
| ONR772 | TTTTTGTCCGTGGAATGAACAATGGAGGGACACGGCGAATAGCCA |
| ONR773 | ATCTATCAAAGGGGACCTCC |
| ONR774 | ACCTGCTGAGTGGATAAAGG |
| ONR775 | TCGCATCCAAGTCTGCATAG |
| ONR776 | TAGCCTCTAAGCTCATGCTG |
| ONR777 | CGTCCGTAGATGAACTTAGG |
| ONR778 | AAACGAGTCCGAGAGAAACG |
| ONR779 | CATCTACATATCACAACGTGCGTGGCTCGATGGCTGTAAGTATCC |
| ONR780 | TTAATGGATTGCTGACCACTTCACCCCACGCACGTTGTGATATGT |
| ONR781 | TGGTAACTGCTGGCTAATACCATGGCCATTGTTCATTCCACGGAC |
| ONR782 | TTTTTGTCCGTGGAATGAACAATGGCCATGGTATTAGCCAGCAGT |
| ONR783 | CGGCAATAACCGTATTTGTC |
| ONR784 | CTACGGCTGGAACAAGAAAC |
| ONR785 | GATAGCACCATTTGCGATAG |
| ONR786 | GCAATGGAGTGTCATTCATC |
| ONR787 | CTGATAATCGAAGCCTTGTG |
| ONR788 | GTCTCGATTCACTGTTAACG |
| ONR789 | TTGAAGTGACCCGGAAATAC |
| ONR790 | GGCGATAGACAGTGATAACC |
| ONR791 | CATCTACATATCACAACGTGCGTGGGGTGAAGTGGTCAGCAATCC |
| ONR593 | GGTCTGACAGTTACCAATGC |
| ONR594 | TCGTTCCACTGAGCGTCAGA |
| ONR597 | GCATTGGTAACTGTCAGACCCCACGCACGTTGTGATATGT |
| ONR598 | TCTGACGCTCAGTGGAACGACCATTGTTCATTCCACGGAC |
| ONR705 | GCCGTCCTGATAGGCTTTGATGATCGGCATTGACCCTGAGTGATT |
| ONR706 | GACCACTACCGCTCTTTTGTCGTTTGGGCCTCGTGATACGCCTAT |
| ONR707 | TAAAAATAGGCGTATCACGAGGCCCAAACGACAAAAGAGCGGTAG |
| ONR651 | CATAAGCCGTAATCACCAGG |
| ONR652 | AGGATTTATCGGGCCAGTTG |
| ONR653 | TAATTCCAGTTTAAAAGGAGGTGAAAACCTCCTTTTTATGACATCCAACA |
| ONR550 | TGTTGGATGTCATAAAAAGGAGGTTTTCACCTCCTTTTAAACTGGAATTACATATGAATATCCTCCTTAG |
| ONR549 | GGGCTGGGGAGGCGGCGCTTTGTTGCTAAATGATCTGGTTTAAAATGGATGTGTAGGCTGGAGCTGCTTC |
| ONR654 | ATCCATTTTAAACCAGATCATTTAGCAACAAAGCGCCGCCTCCCCAGCCC |
| ONR655 | CGGAGTGGAATTCGATTTTG |
| ONR656 | TAAGGTATACCCGGAAGGTG |
| ONR708 | AGAAAAATCACTCAGGGTCAATGCCGATCATCAAAGCCTATCAGG |

Underlines indicate the additional overhangs.

**Table S2: Primer sets and templates used for PCR**

| **DNA fragment** | **Size (bp)** | **Forward primer** | **Reverse primer** | **Template** |
| --- | --- | --- | --- | --- |
| λ_1 | 9757 | ONR454 | ONR178 | λ cI857 genome |
| λ_2 | 9773 | ONR179 | ONR180 | λ cI857 genome |
| λ_3 | 9742 | ONR709 | ONR182 | λ cI857 genome |
| λ_4 | 9727 | ONR183 | ONR184 | λ cI857 genome |
| λ_5 | 9754 | ONR185 | ONR455 | λ cI857 genome |
| λ_3(Δea47)A | 3387 | ONR181 | ONR513 | λ cI857 genome |
| λ_3(Δea47)B | 4716 | ONR514 | ONR182 | λ cI857 genome |
| λ_3(Δea31-ea59)A | 5092 | ONR181 | ONR515 | λ cI857 genome |
| λ_3(Δea31-ea59)B | 1758 | ONR516 | ONR182 | λ cI857 genome |
| λ_3(Δp35-orf61) | 9766 | ONR181 | ONR519 | λ cI857 genome |
| λ_4(Δp35-orf61)B | 7860 | ONR520 | ONR184 | λ cI857 genome |
| λ_4(Δorf61-gam)A | 1939 | ONR183 | ONR517 | λ cI857 genome |
| λ_4(Δorf61-gam)B | 5560 | ONR518 | ONR184 | λ cI857 genome |
| λ_4(Δexo)A | 1914 | ONR183 | ONR522 | λ cI857 genome |
| λ_4(Δexo)B | 6750 | ONR521 | ONR184 | λ cI857 genome |
| λ_4(Δbet)A | 2998 | ONR183 | ONR526 | λ cI857 genome |
| λ_4(Δbet)B | 5980 | ONR525 | ONR184 | λ cI857 genome |
| λ_4(Δkil-sieB)A | 4237 | ONR183 | ONR645 | λ cI857 genome |
| λ_4(Δkil-sieB)B | 3785 | ONR646 | ONR184 | λ cI857 genome |
| λ_4(ΔrexB-cI)A | 6450 | ONR183 | ONR647 | λ cI857 genome |
| λ_4(ΔrexB-cI)B | 968 | ONR648 | ONR184 | λ cI857 genome |
| λ_5(Δren-ninI)A | 1636 | ONR185 | ONR611 | λ cI857 genome |
| λ_5(Δren-ninI)B | 4668 | ONR610 | ONR455 | λ cI857 genome |
| λ_5(Δbor-p79)A | 7852 | ONR185 | ONR506 | λ cI857 genome |
| λ_1(Δbor-p79) | 9782 | ONR505 | ONR178 | λ cI857 genome |
| λ_5(Δnin Δbor)B | 2791 | ONR610 | ONR455 | λ cI857 Δbor-p79 genome |
| λ_4(Δkil, ΔcI) | 4880 | ONR183 | ONR647 | λ cI857 Δkil-seiB genome |
| λ_5(ΔcI Δnin Δbor) | 5179 | ONR648 | ONR455 | λ cI857 Δnin Δbor genome |
| cos(T1) | 345 | ONR748 | ONR749 | λ cI857 genome |
| T1_1 | 9819 | ONR750 | ONR751 | T1 phage genome |
| T1_2 | 9827 | ONR752 | ONR753 | T1 phage genome |
| T1_3 | 9710 | ONR754 | ONR755 | T1 phage genome |
| T1_4 | 9958 | ONR756 | ONR757 | T1 phage genome |
| T1_5 | 9782 | ONR758 | ONR759 | T1 phage genome |
| cos(T3) | 317 | ONR760 | ONR761 | λ cI857 genome |
| T3_1 | 9439 | ONR762 | ONR763 | T3 phage genome |
| T3_2 | 9493 | ONR764 | ONR765 | T3 phage genome |
| T3_3 | 9459 | ONR766 | ONR767 | T3 phage genome |
| T3_4 | 10018 | ONR768 | ONR769 | T3 phage genome |
| cos(T7) | 317 | ONR770 | ONR771 | λ cI857 genome |
| T7_1 | 10203 | ONR772 | ONR773 | T7 phage genome |
| T7_2 | 9864 | ONR774 | ONR775 | T7 phage genome |
| T7_3 | 10114 | ONR776 | ONR777 | T7 phage genome |
| T7_4 | 9797 | ONR778 | ONR779 | T7 phage genome |
| cos(φ80) | 317 | ONR780 | ONR781 | λ cI857 genome |
| φ80_1 | 9099 | ONR782 | ONR783 | φ80 genome |
| φ80_2 | 9288 | ONR784 | ONR785 | φ80 genome |
| φ80_3 | 9362 | ONR786 | ONR787 | φ80 genome |
| φ80_4 | 9595 | ONR788 | ONR789 | φ80 genome |
| φ80_5 | 9044 | ONR790 | ONR791 | φ80 genome |
| pBR322(cos) | 4248 | ONR593 | ONR594 | pBR322 |
| cos(pBR322) | 308 | ONR597 | ONR598 | λ cI857 genome |
| pBRcos(48k) | 2819 | ONR705 | ONR706 | pBR322-cos |
| P1_1 | 13139 | ONR707 | ONR651 | P1 phage genome |
| P1_2 | 13532 | ONR652 | ONR653 | P1 phage genome |
| cat fragment | 1116 | ONR550 | ONR549 | pKD3 |
| P1_3 | 9115 | ONR654 | ONR655 | P1 phage genome |
| P1_4 | 8520 | ONR656 | ONR708 | P1 phage genome |

**Table S3: Combination of PCR fragments**

| **Name** | **Fragment 1** | **Fragment 2** | **Fragment 3** | **Fragment 4** | **Fragment 5** | **Fragment 6** |
| --- | --- | --- | --- | --- | --- | --- |
| **λ　phage** | λ_1 | λ_2 | λ_3 | λ_4 | λ_5 |  |
| **λ(Δea47)** | λ_1 | λ_2 | λ_3(Δea47)A | λ_3(Δea47)B | λ_4 | λ_5 |
| **λ(Δea31-ea59)** | λ_1 | λ_2 | λ_3(Δea31-ea59)A | λ_3(Δea31-ea59)B | λ_4 | λ_5 |
| **λ(Δp35-orf61)** | λ_1 | λ_2 | λ_3(Δp35-orf61) | λ_4(Δp35-orf61)B | λ_5 |  |
| **λ(Δorf61-gam)** | λ_1 | λ_2 | λ_3 | λ_4(Δorf61-gam)A | λ_4(Δorf61-gam)B | λ_5 |
| **λ(Δexo)** | λ_1 | λ_2 | λ_3 | λ_4(Δexo)A | λ_4(Δexo)B | λ_5 |
| **λ(Δbet)** | λ_1 | λ_2 | λ_3 | λ_4(Δbet)A | λ_4(Δbet)B | λ_5 |
| **λ(Δkil-sieB)** | λ_1 | λ_2 | λ_3 | λ_4(Δkil-sieB)A | λ_4(Δkil-sieB)B | λ_5 |
| **λ(ΔrexB-cI)** | λ_1 | λ_2 | λ_3 | λ_4(ΔrexB-cI)A | λ_4(ΔrexB-cI)B | λ_5 |
| **λ(Δren-ninI)** | λ_1 | λ_2 | λ_3 | λ_4 | λ_5(Δren-ninI)A | λ_5(Δren-ninI)B |
| **λ(Δbor-p79)** | λ_1(Δbor-p79) | λ_2 | λ_3 | λ_4 | λ_5(Δbor-p79)A |  |
| **λ(Δnin Δbor)** | λ_1 | λ_2 | λ_3 | λ_4 | λ_5(Δren-ninI)A | λ_5(Δnin Δbor)B |
| **λ(Δkil ΔｃI Δnin Δbor)** | λ_1 | λ_2 | λ_3 | λ_4(Δkil, ΔcI) | λ_5(ΔcI Δnin Δbor) |  |
| **T1(+cos)** | cos(T1) | T1_1 | T1_2 | T1_3 | T1_4 | T1_5 |
| **T3(+cos)** | cos(T3) | T3_1 | T3_2 | T3_3 | T3_4 |  |
| **T7(+cos)** | cos(T7) | T7_1 | T7_2 | T7_3 | T7_4 |  |
| **φ80(+cos)** | cos(φ80) | φ80_1 | φ80_2 | φ80_3 | φ80_4 | φ80_5 |
| **pBRcos-48k** | pBRcos(48k) | P1_1 | P1_2 | cat fragment | P1_3 | P1_4 |
| **22 kb for Exo III assembly (for Fig. S1)** | λ(4.6k) | λ(0.3k) | λ(4.8k) | P1(3.3k) | P1(3.0k) | P1(6.0k) |

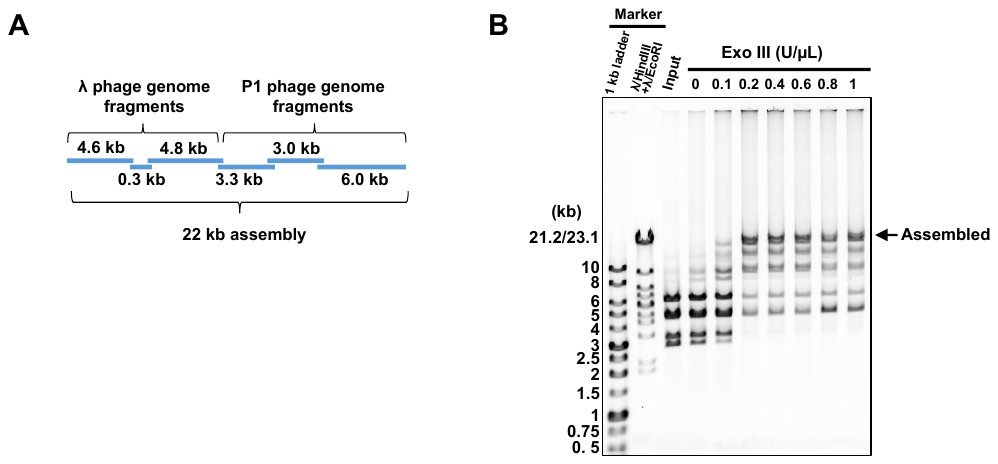

**Figure S1.** Examination of DNA assembly conditions by using Exo III

A. DNA fragments used for examination of assembly conditions. PCR fragments of the indicated sizes prepared from theλ phage genome and the P1 phage genome as templates were used for Exo III assembly. Adjacent fragments overlap by 50 bp each.

B. DNA assembly according to the concentration of Exo III. Exo III assembly was carried out as describe in the meterials and methods section, except that the concentration of Exo III was changed from 0 to 1 mU as indicated. After assembly reaction, the assembled DNA was analysed by 0.7% agarose gel electrophoresis. Arrow indicates the assembled DNA.

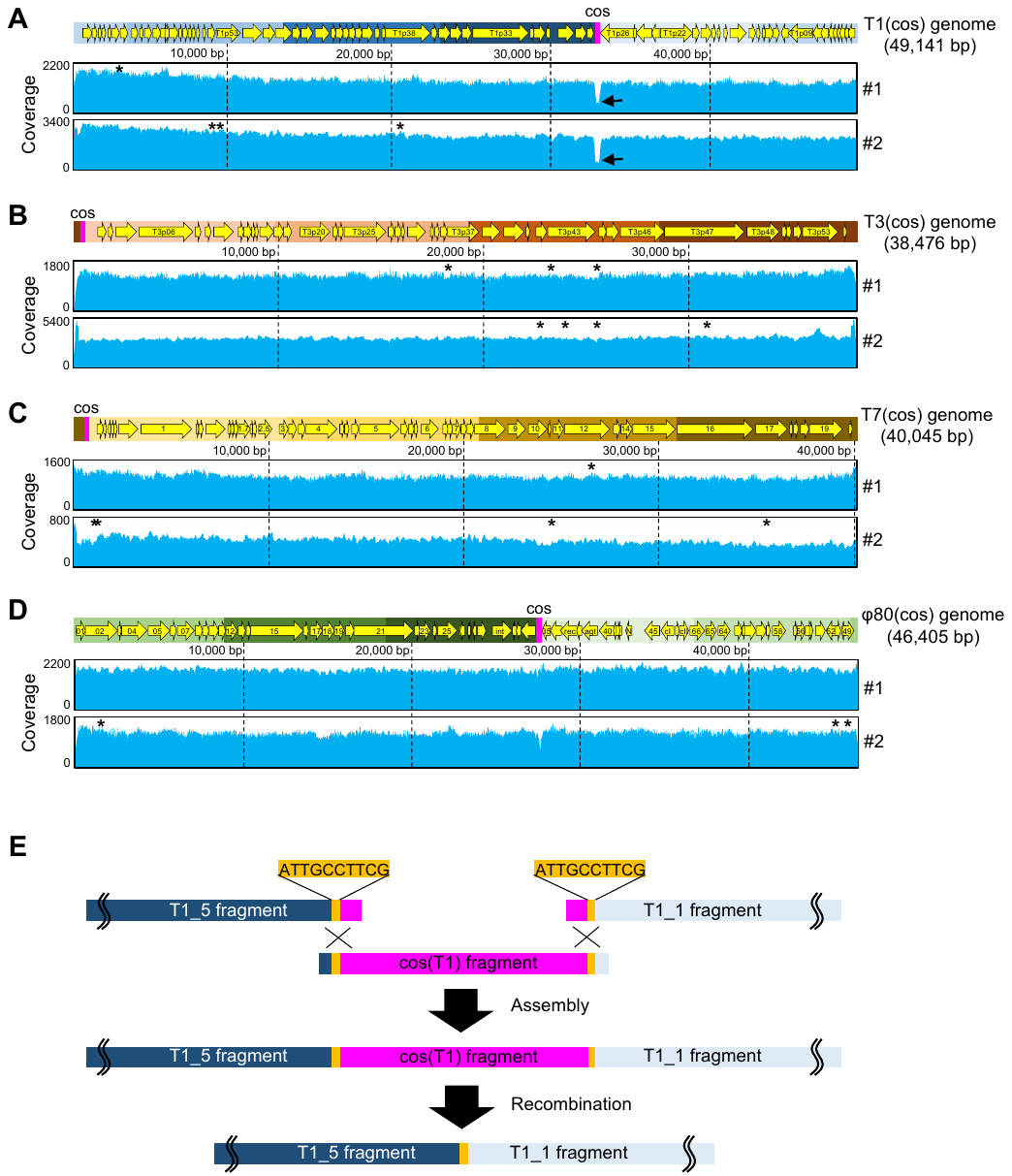

**Figure S2**

NGS analysis of various phages constructed by iPac.

A - D. Genomes of constructed T1 phage (A), T3 phage (B), T7 phage (C) and φ80 (D) were analyzed by NGS. The reads obtained by NGS were mapped to the reference genome sequences. Asterisks indicate single nucleotide substitution mutations. Arrows indicate the drop of reads in the inserted cos region.

E. Expected mechanism of cos deletion in T1 phage construction. 10 bp homologous sequences that occurred during the fragments design step are indicated in orange and cos fragment in magenta.
